# Subsurface Zircons with Presumptive “Biogenic” Inclusions as Potentially Useful Proxies for Studying Precambrian Bygone Biospheres in Goa

**DOI:** 10.1101/210013

**Authors:** Dabolkar Sujata, Kamat Nandkumar

## Abstract

This work was inspired by recent report by Bell et al., 2015 who studied potentially biogenic carbon preserved in a 4.1 billion-year-old Zircon and need to assess the potential of Zircons found in Goa. Zircons (ZrSiO4) are naturally occurring silicate minerals which show radioactivity and high ductility and contain traces of Thorium and Uranium useful in Uranium–Thorium /Thorium −230 dating techniques. Zircons can be found in igneous, metamorphic rocks, sedimentary deposits and occurs as a detrital minerals in river and beach sands. Previous reports show that the Zircons can occur in different shapes such as round, elongated and with surface characteristics (Gartner et al.,2013). U-Pb Zircon dating methods had been used to study the continental growth in the western Dharwar craton of southern India (Jayananda et al., 2015). The present study was aimed at detection of subsurface Zircons with biogenic inclusions and assess their use as proxies for studying bygone Precambrium biospheres in Goa. Deep tubewell drilled Cores (60 and 65 m deep from surface) in island of Tiswadi at Taleigao were analyzed by light microscopy, Phase contrast microscopy and SEM to detect and classify the Zircons. In rapid preliminary sampling, total 50 Zircons were identified and 98% indicated the presence of interesting inclusions. These could be bubbles or kerogens or unidentified biological material. Zircons were classified as elongated, slightly rounded with sharp edges and showed widespread variety of surface characteristics like fracturing, cracks, scratches, striations and impact pits which may occur during transport processes. It is suggested that Zircons with presumptive biogenic inclusions can be further studied using techniques such as Raman Spectroscopy, Carbon Isotopic Measurements, X-Ray Microscopy, Trace Element Measurement consistent with Bell et al., 2015. More exhaustive studies have been undertaken to create a detail image database of Zircons from various other local samples to pinpoint those specifically useful for advanced work based on image analysis of the presumptive bioinclusions. Further attempts would be made to develop specific harvesting techniques to select potentially useful Zircons. International collaborations would be sought for applications of advanced techniques to local Zircons. Such studies would shed light on nature of bygone Precambrian biospheres in Goa and help in understanding evolution of life and the impact of plate tectonics and cataclysmic events shaping life on this planet.

## Introduction

The aim of this study was detection of subsurface Zircons with biogenic inclusions and assess their use as proxies for studying bygone Precambrian biospheres in Goa. Zircons has played a prominent and complex role in interpreting the composition and history of modern and ancient sediments. Presence of carbon in 4.1 billion year zircon was studied by Bell et al., 2015. During this work efforts were made to separate, classify and carry out microscopic studies of the Zircons obtained from the deep tubewell drilled Cores (60 and 65 m deep from surface) in island of Tiswadi at Taleigao. SEM studies of the zircons were carried out. Such studies would shed light on nature of bygone Precambrian biospheres in Goa (Fig 2) and help in understanding evolution of life and the impact of plate tectonics and cataclysmic events shaping life on this planet.

## Materials and Methods

### Regional geologic setting

Goa is situated in the north western part of the metallogenic archean Western Dharwar Craton. The Dharwar Craton is divided into Eastern and Western Cratons wherein Goa is situated in the north western part of the WDC which includes Sanvordem, Bicholim, and Vagheri Formations (Dessai 2010).Tiswadi island is a part of Sanvordem formation constituting the metagreywacke with subordinate metaconglomerate, lensoid tilloid samples (Dessai 2011).

Deep tube well drilled Cores (60 and 65 m deep from surface) in island of Tiswadi at Taleigao were obtained from A.G Chachadi, identified as lensoid tilloid (Fig 2). Samples were powdered as shown in figure 2a and figure 2b, sieved and subjected to washing. Direct DPX mount, Scanning electron microscopy (SEM) and Light and phase contrast microscopic studies were carried out. 24bitmapped Images processed using SCION software(4.0.2) for following parameters.1.Find edge function output, 2. The density slice function output, and 3. The surface pixel plot density (SPPD).

## Results

Both 60m and 65m deep core samples showed high fraction of Zircons in preliminary sampling. Total 50 zircons were identified and 98% of Zircons indicated the presence of interesting inclusions. The sieving and floatation technique helps in enriching the fractions with zircons, which can be directly observed under light microscopy (fig 3). The captured images of zircons were imported and converted to 24 bitmapped images using SCION image processing software (USA) beta, freeware version 4.0.2 (an image processing and analysis program for the IBM PC) to get distinct image panels for each Zircon with respective DIA output-original image, find edge function (FEF), and surface pixel plot density (SPPD). These panels are shown in Figure 4 and 5. Microscopic techniques helped in the study of presence of presumptive bio inclusions inside the zircon as shown in the figure 6a to 6d.

**Fig 1a:**
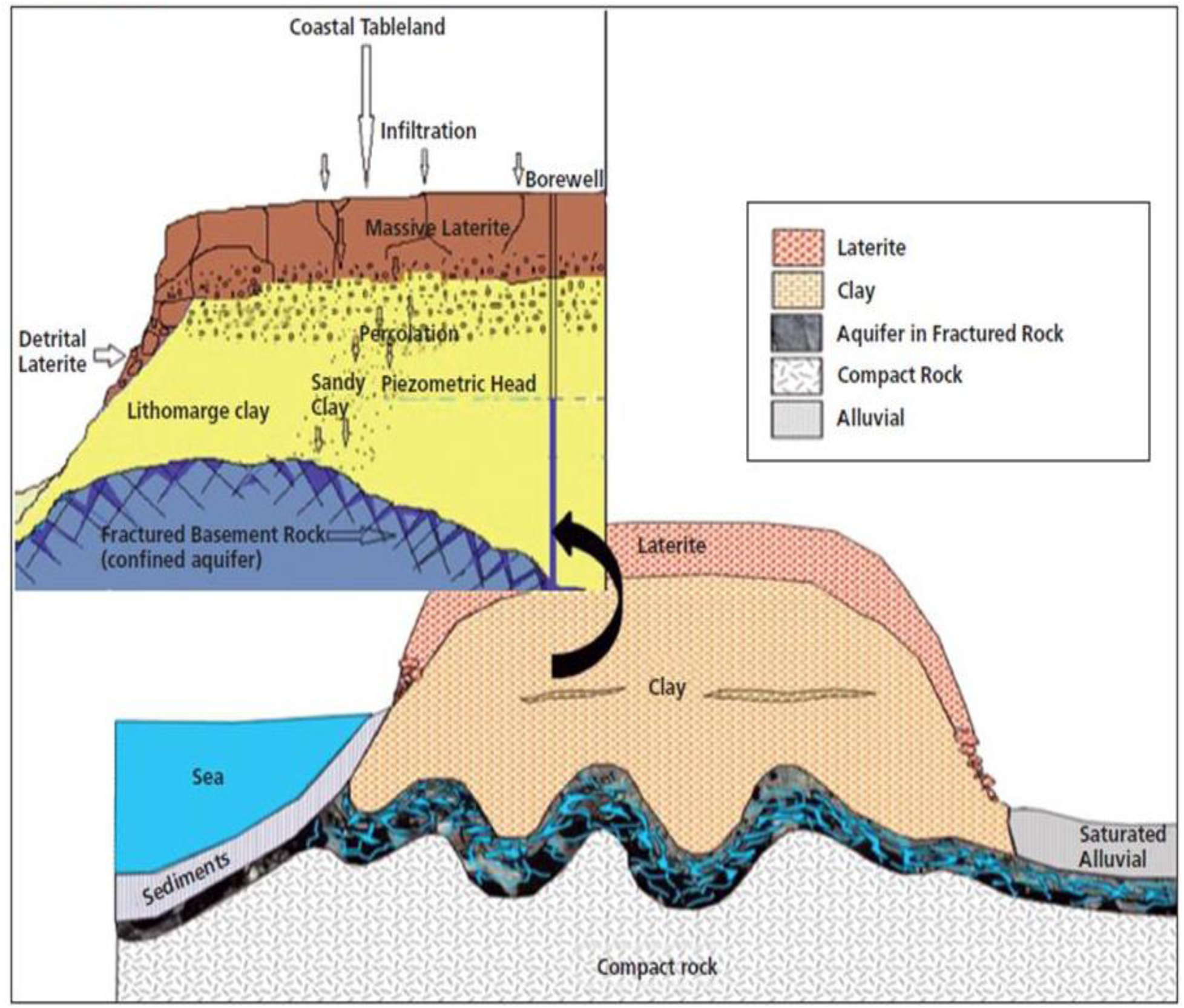
Geohydrological setting of tubewells drilled (Chachadi)

**Fig 1b:**
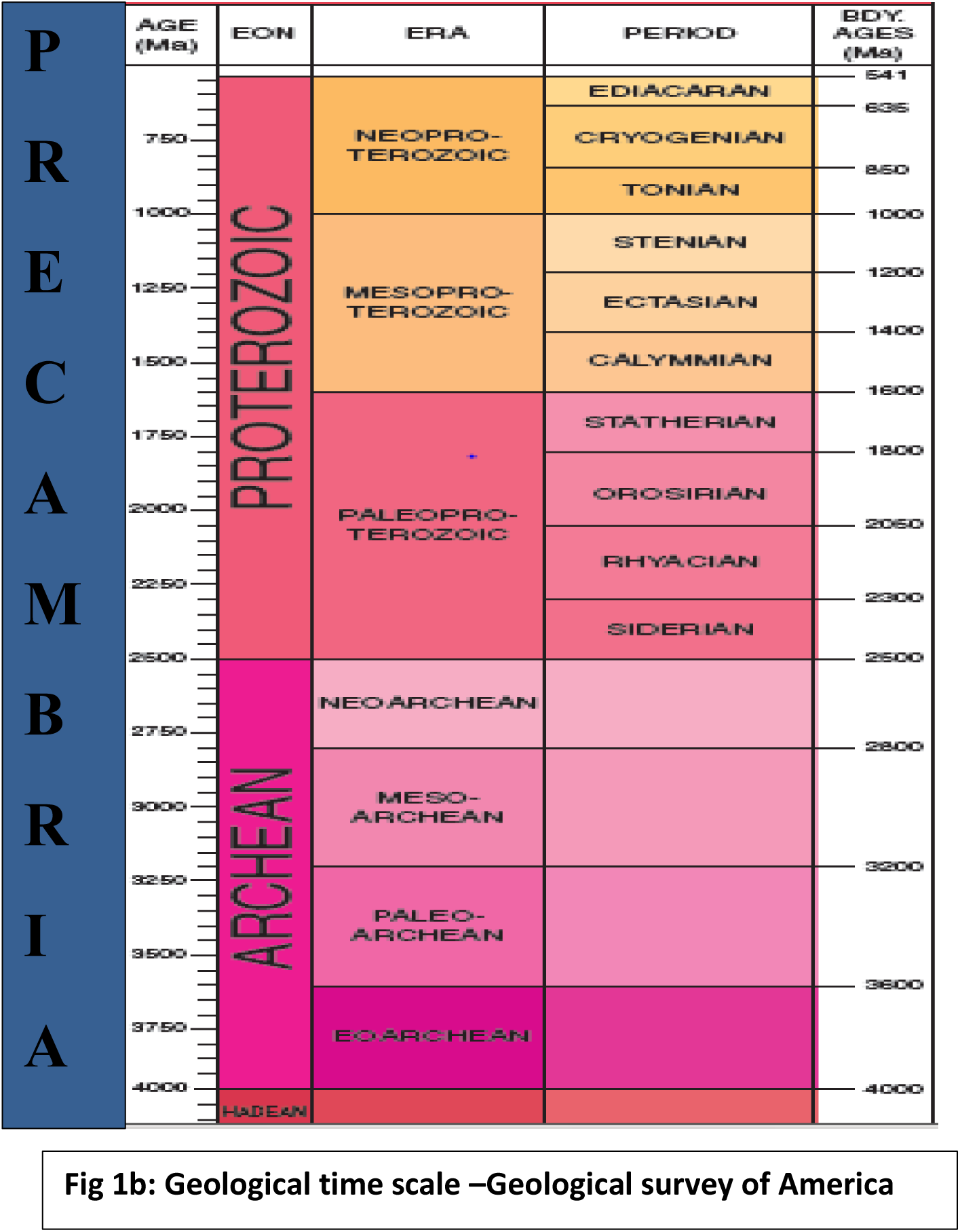
Geological time scale–Geological survey of America

**Figure 2:**
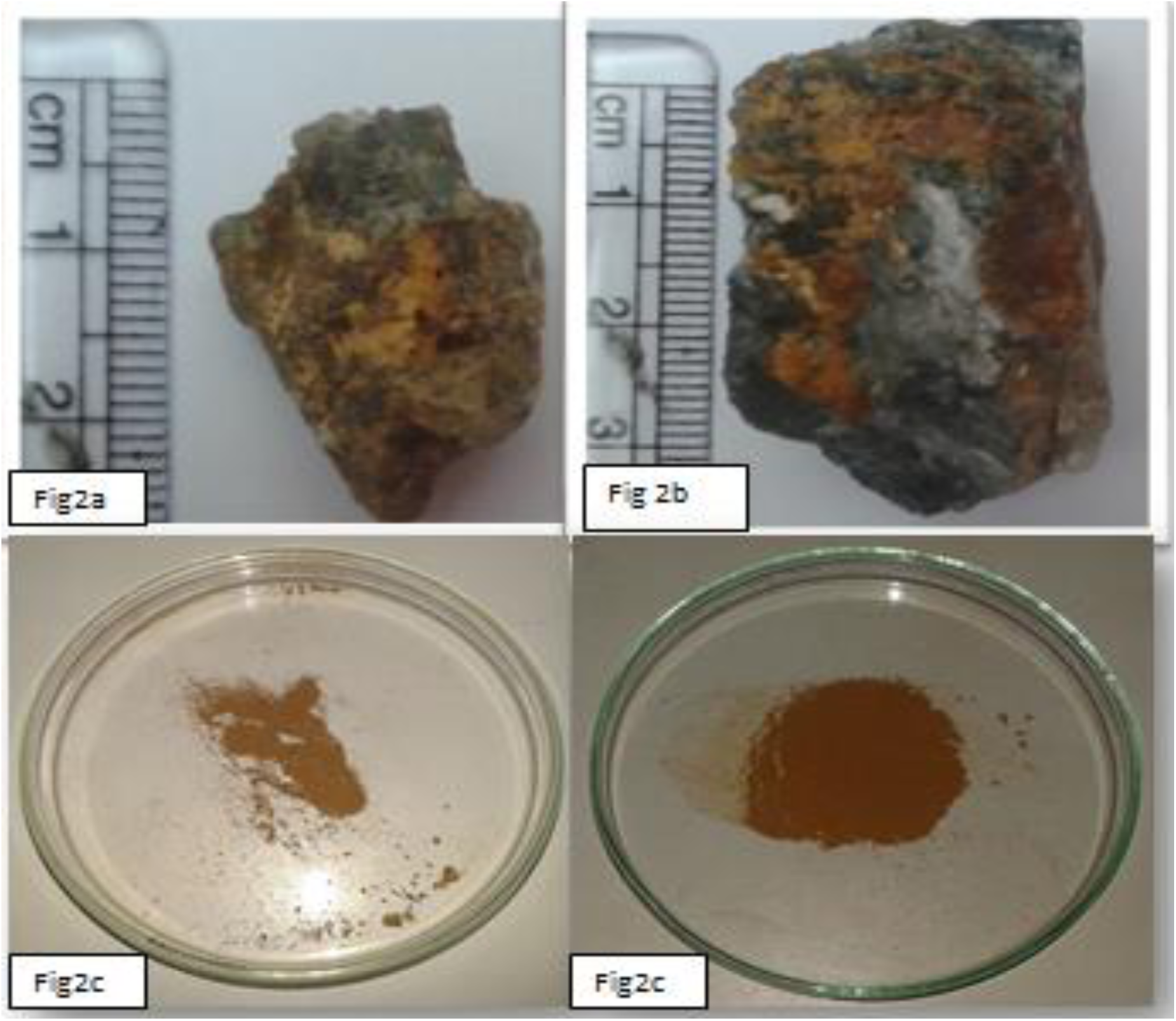
Lensoid tilloid samples Powdered tilloid samples

**Figure 3:**
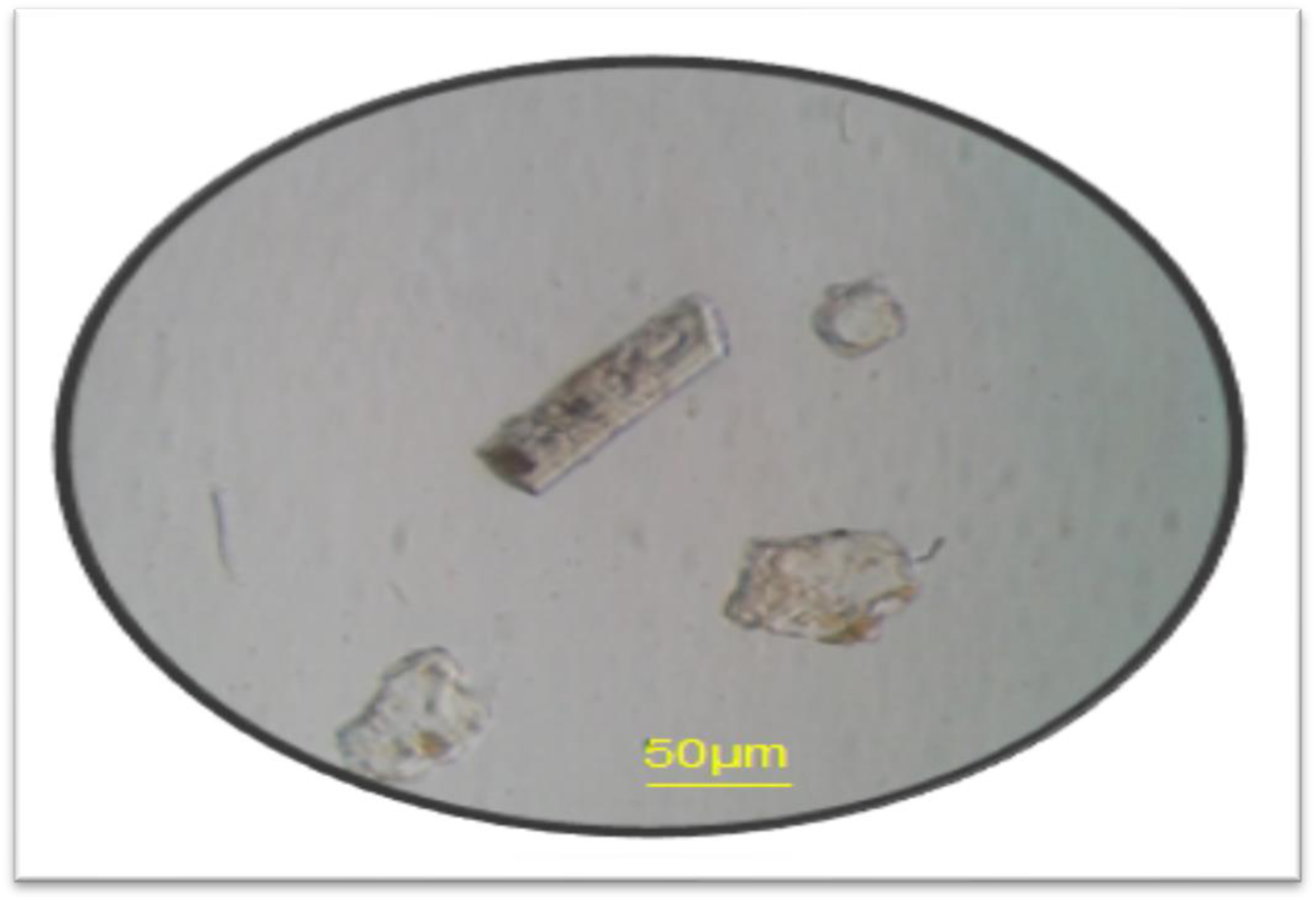
Mixed field showing the Zircon and other minerals

**Fig 4:**
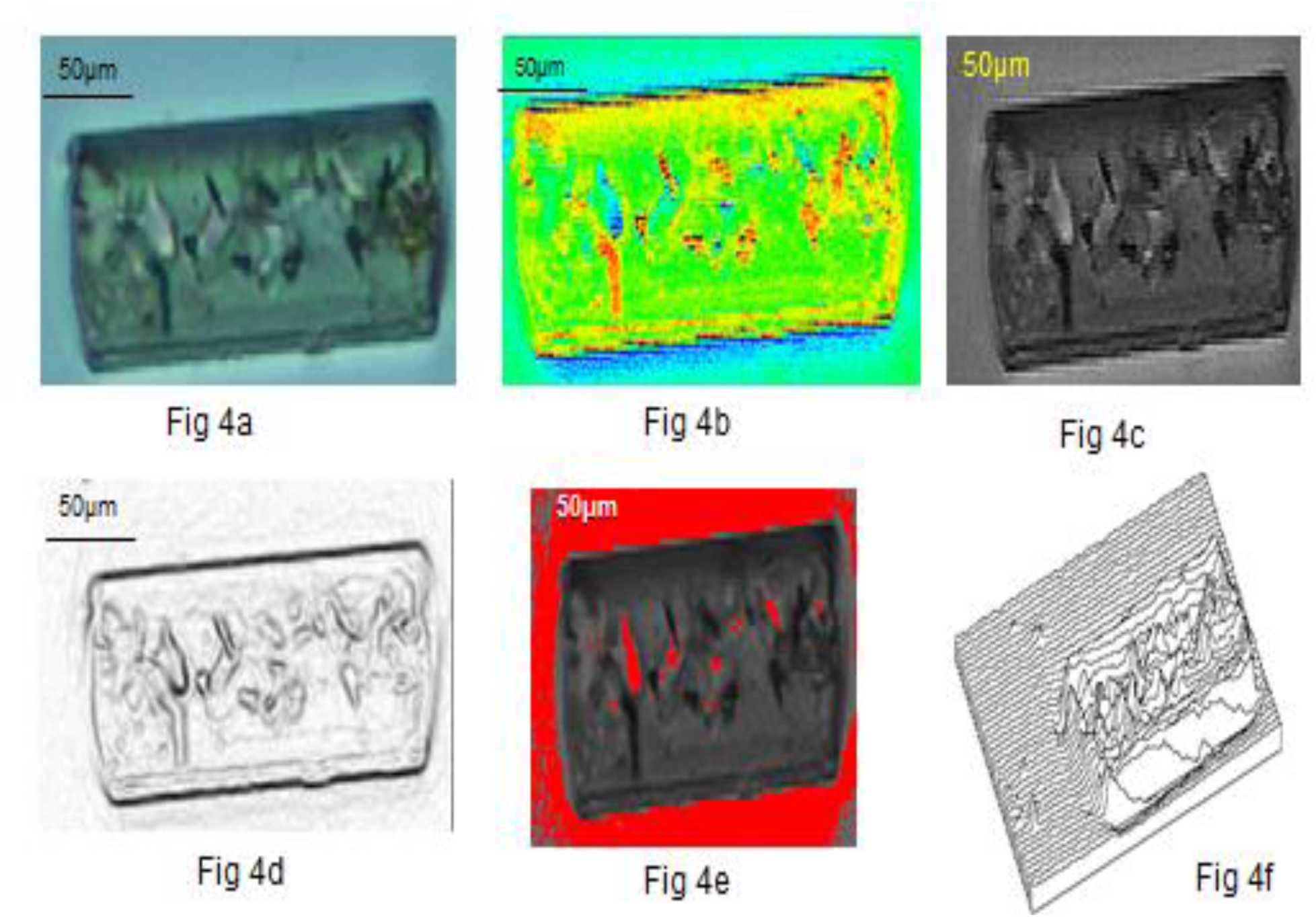
original Zircon, 4b-Pseudo, 4c -sharp edges of zircon, 4d-sharp edges of zircons, 4e-density slice, 4f -surface plot

**Figure 5:**
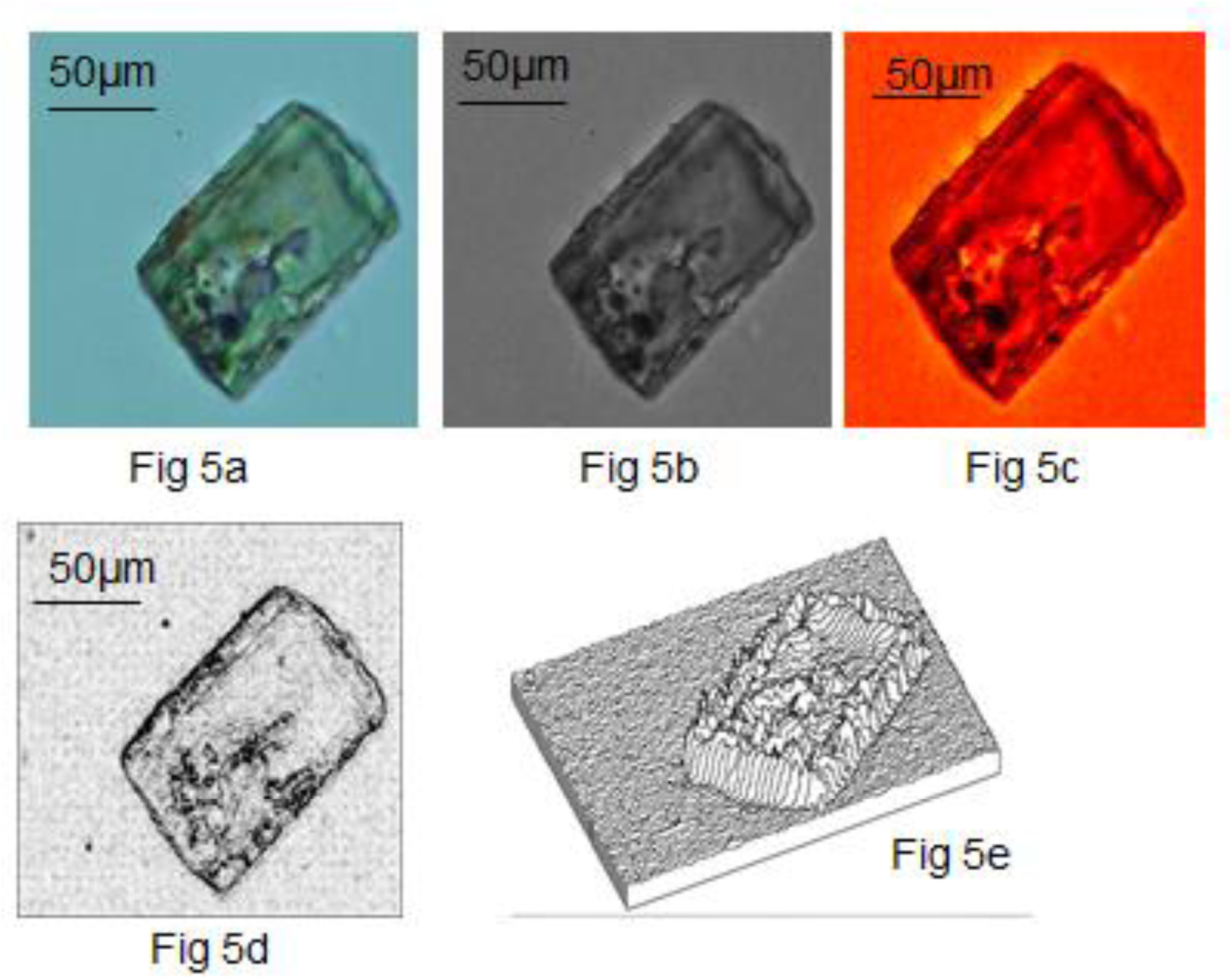
a-original Zircon, b-sharp edges of zircon, c -Pseudo, d-sharp edges of zircons and e-surface plot

**Fig6:**
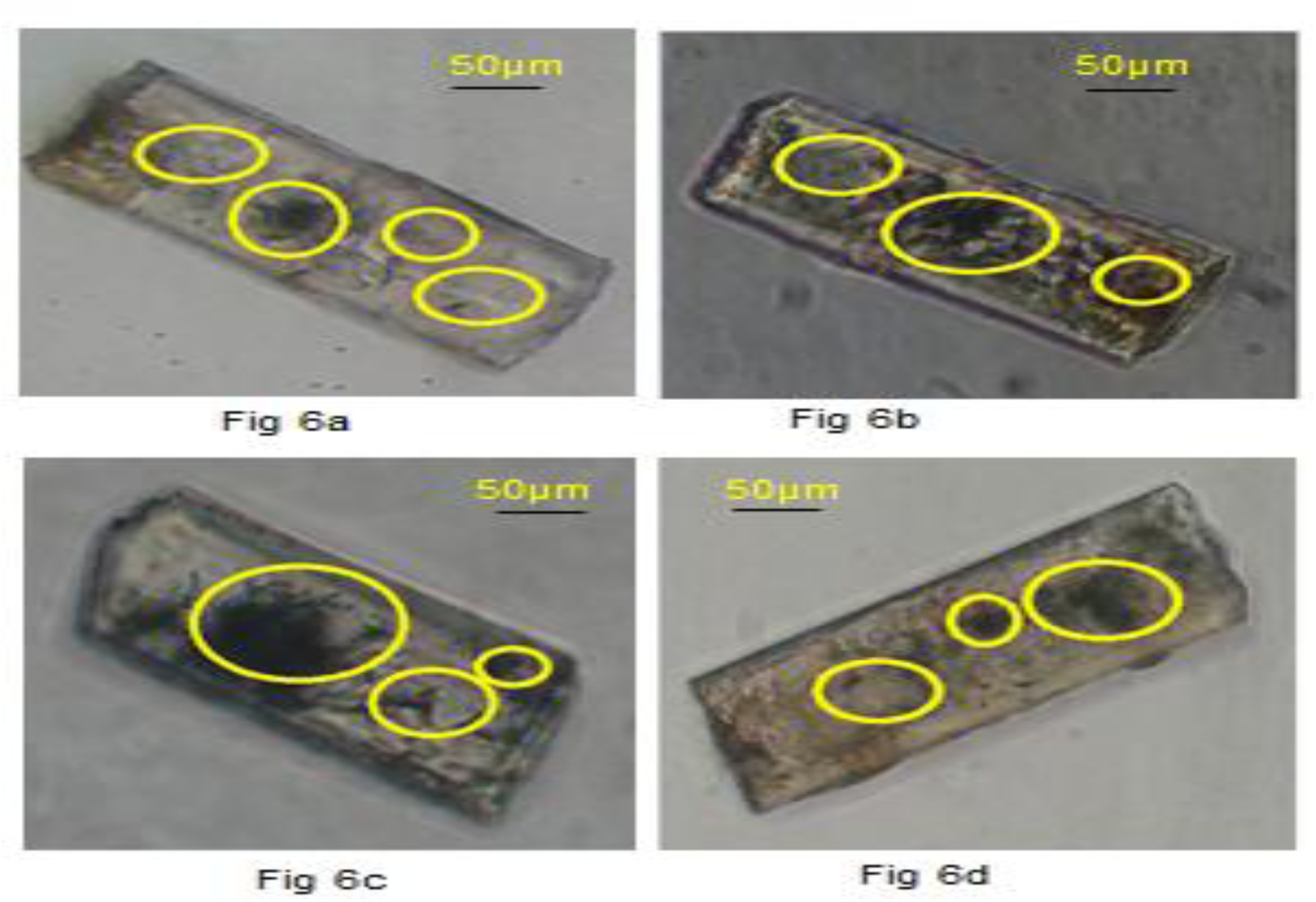
Yellow circles indicate presumptive bioinclusions

**Figure 7:**
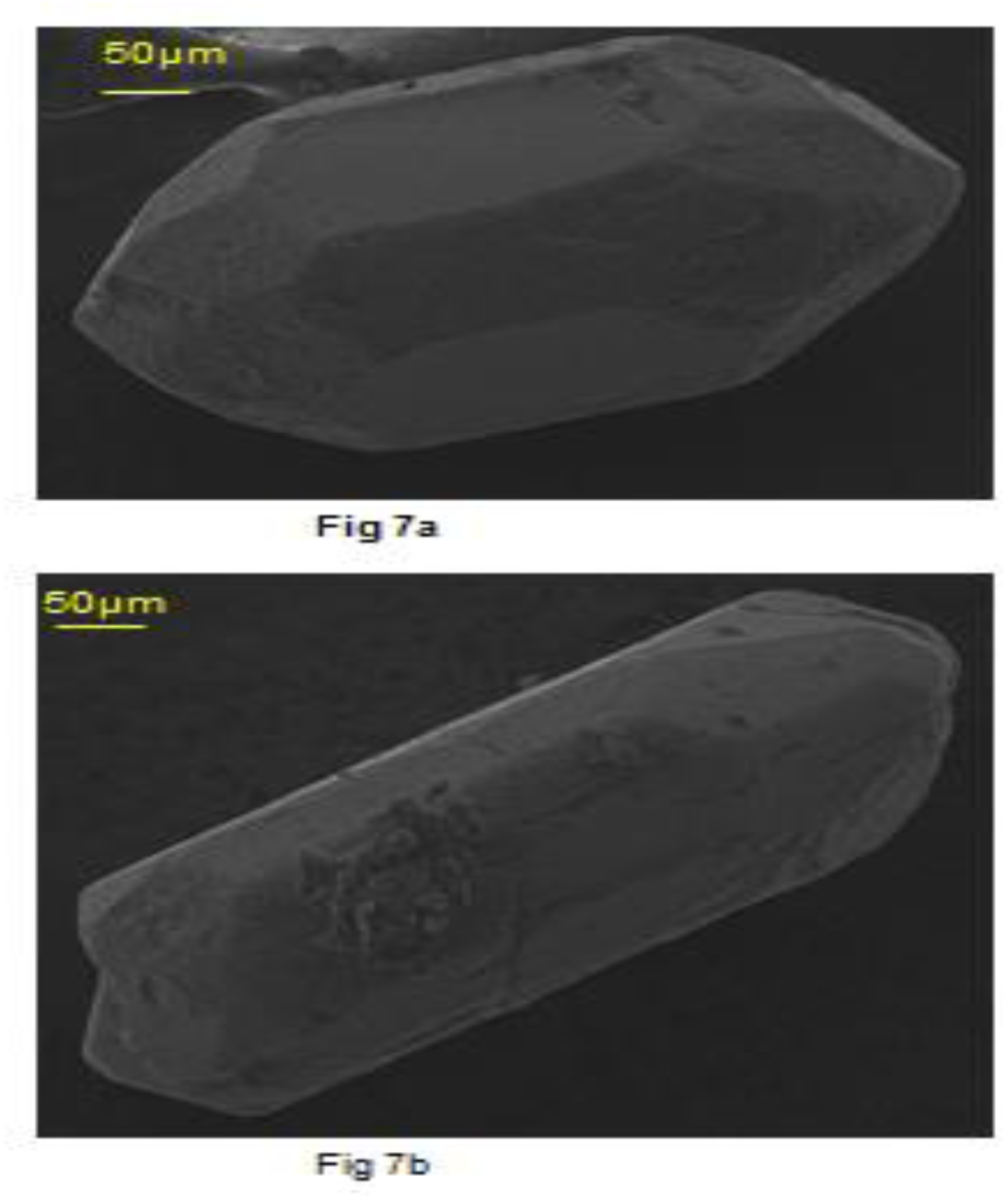
SEM typology of Zircon

## Discussion

The results show that using laboratory techniques and advanced image analysis software it is possible to visualize the Zircons and bioinclusions. It is suggested that zircons with presumptive biogenic inclusions can be further studied using techniques such as Raman Spectroscopy, Carbon Isotopic Measurements, X-Ray Microscopy, Trace Element Measurement consistent with Bell et al., 2015. More exhaustive studies have been undertaken to create a detail image database of Zircons from various other local samples to pinpoint those specifically useful for advanced work based on image analysis of the presumptive bioinclusions. Further attempts would be made to develop specific harvesting techniques to select potentially useful Zircons. International collaborations would be sought for applications of advanced techniques to local zircons. Such studies would shed light on nature of bygone Precambrian biospheres in Goa and help in understanding evolution of life and the impact of plate tectonics and cataclysmic events shaping life on this planet(Bell et., 2015).

## Acknowledgements

We thank Anne Berger, Sales Manager, Digital Surf, France for giving permission to use Mountains Map software, for SEM image processing and analysis. This work was supported by UGC-SAP Phase II – Biodiversity, Bioprospecting programme and Goa University Fungus Culture Collection (GUFCC). We thank R.N.S Bandekar CO, Vasco da Gama for funding the work on biomineral studies and Professor A.G Chachadi from department of Earth Science for proving core samples.

### Figure captions

Figure 1a: Geohydrological setting of tubewells drilled (Chachadi, 2013)

Figure 1b: Geological time scale

Figure 2 (a-b): Lensoid tilloid samples

Figure 2 (c-d): Powdered tilloid samples

Figure 3: Mixed field showing the Zircon and other minerals

Figure 4(a-f): original Zircon, 4b- Pseudo, 4c -sharp edges of zircon, 4d-sharp edges of zircons, 4e-density slice, 4f -surface plot

Figure 5(a-e): a-original Zircon, b- sharp edges of zircon, c -Pseudo, d-sharp edges of zircons and e-surface plot

Figure 6 (a-b): Yellow circles indicate presumptive bioinclusions

Figure 7 (a-b): SEM typology of Zircon

